# Pick your battles: competitive strategies of green chromides (*Etroplus suratensis)* during inter- and intraspecific contests

**DOI:** 10.64898/2026.10.08.757801

**Authors:** Vandana Pradeep, Pranav Kulkarni, Ratna Ghosal

## Abstract

Competitive ability is a key determinant of organismal fitness, yet high costs of interference competition necessitate strategic behavioural responses. This study investigated how green chromide (GC), a cichlid native to the Indian subcontinent, competes with the globally invasive tilapia species (T, *Oreochromis* spp.) that has encroached upon its habitats. We staged dyadic feeding competitions between GC and T, and analysed aggression, feeding, and hiding behaviours of both the species and further contrasted them with intraspecific contests in GC. We varied developmental stages, residency status of opponents, and used different combinations of body sizes of competitors to test their effects on the outcome of dyadic contests. We hypothesized that competitors would adjust their strategies based on the context of the competition, and thus, we aimed to identify the factors that contributed to such decision-making. Our findings showed residency as the primary driver of competitive strategy, wherein residents had higher aggression and feeding when compared to intruders. The developmental stage had a significant effect on feeding, increasing from young to older stages, but body size contrasts did not contribute significantly towards the outcome of the contests. For the species effect, T emerged as a winner in most of the interspecific contests, exhibiting a significant amount of total aggression. However, GC showed distinct behavioural strategies when competing against T versus conspecifics, and exhibited higher levels of aggression and reduced hiding during intra- when compared to interspecific trials. Surprisingly, the feeding behaviour of GC was not significantly impacted in the presence of T, possibly because GC used hiding as a defence strategy, emphasizing the context-dependent, strategic thinking of the species. Overall, the variety of behavioural tactics exhibited by GC in staged competitive interactions showcases robust survival skills of the species, a key trait that may help maintain its fitness in challenging ecological habitats.

**Highlights:**

- We tested competitive strategies of cichlids in staged dyadic contests
- Residency significantly influenced the contest outcomes
- Tilapia outcompeted green chromides, particularly, with regards to aggression
- Green chromides maintained feeding in presence of tilapia, using hiding to defend
- Competitors’ capabilities, and not the contest format, dictated the strategies used.

## 1. Introduction

Since Ernst Haeckel defined ecology in 1866 as the study of an animal’s comprehensive relationship with its environment (Egerton, 2013), the field has focused on understanding biotic interactions operating across intra- and interspecific scales (Koffel et al., 2021). Both inter- and intraspecific interactions can further be categorized as either cooperative (Barker et al., 2017; Bronstein & Sridhar, 2024; Sachs, 2006) or competitive (Barabás et al., 2016; Birch, 1957). Though competition has the potential to impose negative fitness costs and may even threaten survival (Kanishka et al., 2025), it is a predominant force in ecosystems worldwide, driven by the fixed nature of resource availability (Grether et al., 2017; Peiman & Robinson, 2010). Consequently, these competitive interactions are inherently dynamic and influence the fundamental structure and function of ecosystems across spatial and temporal gradients.

With regard to competitive interactions, direct or interference competition levies a higher cost to the fitness and survival of organisms when compared to indirect or exploitative interactions (Amarasekare, 2002; Case & Gilpin, 1974). This is mainly because interference competition is mediated by costly signals, for example, expression of ornamental traits (Berglund, 1996; Kenyon & Martin, 2023), behavioural demonstration of aggression (Ward et al., 2006; Wojczulanis Jakubas et al., 2015; Linnell & Strand, 2000), release of chemical signals (Hay, 2008; Stockley et al., 2013), and advertisement of acoustic calls (Amorim et al., 2019; Reichert & Gerhardt, 2014), all of which convey information regarding the competitive ability of the competitors. Since the maintenance of these traits is metabolically demanding, organisms frequently rely on diverse indicators (physical cues) or metrics to evaluate opponent strength, which facilitates strategic decision-making and allows them to avoid physiological costs of direct physical combat (Arnott & Elwood, 2009).

Some of the popular assessment metrics used by the organisms are, differences in body sizes (Arnott & Elwood, 2009; Chapin et al., 2019), including the exaggeration of the ornamental traits (Arnott & Elwood, 2009; Berglund, 1996), and residency [defined as the duration for which an individual exhibits largely uninterrupted occupancy of an area (Chapman et al., 2014; Arnott & Elwood, 2009)], and contrast in the frequency or duration of specific behavioural displays, for example, head shaking (Stuart Fox, 2006), vocal displays (Benítez et al., 2017) and aggression (Wilson et al., 2011). Research indicates that the utilization of these cues varies according to the ecological context, highlighting an adaptive cognitive capacity that allows organisms to respond effectively to local competitive pressures, thereby enhancing their overall fitness.

Additionally, understanding how organisms negotiate interference competition, whether by avoiding or engaging in it (Case & Gilpin, 1974), is vital for species persistence, particularly under the resource constraints characteristic of human-altered environments (Kanishka et al., 2025). Building upon this, our research assesses the competitive strategies of a focal species within both intraspecific and interspecific contexts to elucidate the key drivers modulating these strategic responses. For the focal species, we used green chromide (*Etroplus suratensis*) (abbreviated as GC from here onwards), an economically important cichlid (Padmakumar et al., 2012; Renjitkhumar & Roshni, 2026) that has a very high rate of harvest and is native to the brackish and freshwater bodies of the Indian subcontinent (Mallick et al., 2022; Desai and Ghosal, 2024). The native habitats of GC have been invaded by several alien species, including suckermouth catfish (*Pterygoplichthys* spp.) (Singh & Lakra, 2011), African catfish (*Clarias gariepinu*s) (Krishnakumar et al., 2011), and common carp (*Cyprinus carpio*) (Raj et al., 2021). One such alien species that co-occurs with GC (Joshi et al., 2021; Roshni et al., 2021) is tilapia (*Oreochromis* spp.) (abbreviated as T from here onwards), which originates from Africa (Fryer and Iles, 1972). T is known to be invasive across the globe (Russell et al., 2012; Gracida-Juárez et al., 2022), and it is also a highly aggressive species (Barreto et al., 2011; Sanches et al., 2012). Additionally, since both species belong to the same family, Cichlidae, they have overlapping demands for resources, as documented in Lake Vembanad, Kerala, wherein GC and T showed a huge dietary overlap (Roshni et al., 2021). Given this, we staged dyadic contests between GC and T to unravel GC’s strategies for combating the aggressive T, and further contrasted these with intraspecific competition in GC. Across both inter- and intraspecific competitions, we manipulated body size (total length, TL, measured from the tip of the snout to the tip of the caudal fin; Önsoy et al., 2011) and opponents’ residency to determine whether these traits dictate competitive dynamics. This comparative framework aims to reveal how organisms adapt their behavioural strategies to immediate environmental contexts and the degree to which individual traits influence their strategic decision-making.

## 1. Materials and methods

### 2.1 Study animals and housing conditions

GC were obtained from the Hooghly River near Mousuni Island, West Bengal, India (21°41′44.88′′ N, 88°11′8.714′′ E), during July-August 2022 and again during December 2024-January 2025. T were obtained from freshwater systems in Gujarat, India, including the Sabarmati River near Ahmedabad (22°59′51.3492′′ N, 72°33′51.6132′′ E) in December 2022 and Petli Lake, Kheda district (22°35′56.861′′ N, 72°45′26.604′′ E) during December 2024- January 2025. Since both GC (Bindu & Padmakumar, 2020; Christo & Divya, 2025) and T (Bera et al., 2023) are considered commercial food fish in India, both these fish were procured from local fishermen, who used cast nets to capture them. A total of 100 GC and 25 T were taken at a time from the larger population caught by the fishermen and were transported to the fish laboratory at Ahmedabad University for experiments. GC was transported by train with a journey spanning 36-hours due to the distance of the field site from the laboratory, while T was transported by road with a journey of a maximum duration of 2 hours. In both cases the fish were transported in air filled plastic containers with clean water, and it was ensured that the temperature was maintained using a digital thermometer (Thermisto, Delhi, India) and dissolved oxygen levels were checked and maintained using a portable testing kit (Aquasol, Navi Mumbai, India) to minimise stress. However, since GC had a considerably long transport time, 3-4 water changes were given throughout the 36-hour journey at regular intervals to maintain water quality and aeration. All the fish were housed under laboratory conditions for a minimum of one month before conducting any experiments.

In the laboratory, fish were housed in 70-L glass aquaria measuring 52cm*37cm*32cm (length*width*height). Each aquarium was equipped with an internal power filter (Bluepet, Kolkata, India) for adequate aeration and to maintain water quality. Water temperature and photoperiod were maintained at 27±2°C and 14 h light:10 h dark, respectively. Water temperatures were checked daily using a digital thermometer (Thermisto, Delhi, India), and dissolved oxygen was monitored, every alternate day, while using a portable testing kit (Aquasol, Navi Mumbai, India). Water in the tanks was changed every week to ensure adequate water quality. In case of both GC (for all the developmental stages) and T (except the oldest developmental stage), each home tank housed 5-6 individuals of a species of a given developmental stage, and individuals were accordingly returned to the same home tanks once the trials were over in the experimental tanks. However, in the case of the oldest developmental stage of T, each fish was housed separately in a single home tank because we observed increased aggression among the conspecifics. Fish were fed once daily at 5% of their body weight using commercially available feed, Growfin pellets (Growel, Andhra Pradesh, India). A feeding rate of approximately 5% of the body weight per day was selected to provide a restricted feeding regime which is known to support adequate growth and feed utilization in both GC (Parakkandi et al., 2021) and T (Huang et al., 2015). All individuals were inspected for diseases, injuries, and abnormalities, and only healthy fish were selected for the experiments. Prior to the experimental trials, individual fish were weighed using the water-displacement method. A 1-L beaker containing a known volume of water was first weighed using a weighing balance, after which the fish was briefly transferred into the beaker using hand-held net and the beaker was weighed again. Fish weight was calculated as the difference between the two measurements. The total length (TL, Önsoy et al., 2011) was also measured before the experimental trials using a ruler. Based on the TL, individuals were assigned to three nominal size classes (6, 8, and 11 cm). TL (Mean±SEM) of these size classes was 5.63 ± 0.046 cm, 8.46 ± 0.05 cm, and 10.86 ± 0.071 cm, respectively.

Corresponding body masses were 11.64 ± 0.104 g, 19.6 ± 1.14 g, and 54.77 ± 0.302 g for GC, and 8.51 ± 0.11 g, 13.12 ± 0.078 g, and 49.86 ± 0.33 g for T. Accordingly, individuals were referred to as GC6, GC8, and GC11, and T6, T8, and T11. In T, body weight of 0.003-12 g (6 cm in our study) was considered fry, 12-30 g (8 cm in our study) was considered to be fingerling, and 30-200 g (11 cm in our study) was considered to be juvenile/grower (Food and Agriculture Organization [FAO], 1989). In GC, a length of < 7 cm was considered to be fry, and 7-8 cm was considered to be fingerling (Vikas et al., 2020). While there is no literature on the juvenile developmental stage in GC, studies have shown that sexual maturity is attained only after 12 cm (Bindu and Padmakumar, 2020). Thus, in this study, we classified GC11 as juvenile. We selected these three size classes for both the species, GC and T, to represent different developmental stages, and this allowed us to make several size-matched and size-asymmetric combinations.

### 2.2 Experimental Set-up

All experimental trials were conducted in 70-L glass tanks measuring 52cm*37cm*32cm (length*width*height) with a white background marked with an 8×8 cm grid to quantify the speed of different behaviours. Each tank had an internal power filter (Bluepet, Kolkata, India) and was maintained at 27±2°C under a 14:10 h light–dark cycle. Each trial lasted 45 minutes and was recorded using a digital handycam (Panasonic HC-V385). The fish were caught and moved from the home tanks to the experimental tanks using a hand-held net. The trials began with a 15-min acclimatization period, followed by the introduction of food and a 30-min observation period during which behaviours were noted. 2-3 g of food, equivalent to 4-5% body weight for smaller (6-8 cm) individuals, and 2-3% for 11 cm individuals, was introduced at the end of the acclimatization period per trial. The rationing of the food was maintained as per the feeding guidelines and standard practices [4-6% of the body weight per day for fry and fingerlings (5-20 g), respectively, and ∼3% for juveniles (20-100 g), Food and Agriculture Organization of the United Nations; El-Sayed, 2006].

The trials were broadly classified into inter- and intraspecific trials; each of the trial types was further classified into size-matched and size-unmatched trials. We developed a code for all the trial types, and in the interspecific category, the first part of the code denoted the size class (or developmental stage) of GC, while the second part denoted the size class (or developmental stage) of T. In the case of intraspecific trials, codes had combinations of different size classes of GC, and the larger size class was followed by the smaller one for the size-unmatched intraspecific trials.

For each of the trial types, residency was experimentally manipulated. One individual was designated as the resident (referred to as R from here onwards) and introduced into the experimental tank 24 h before the start of the trial, and remained in the same tank for the next four days. The intruder (referred to as I from here onwards) was introduced at the start of each trial and was returned to the housing tank after the observation period. After four days, R was also returned to the housing tanks. After an interval of 48 h, residency roles were reversed for each pair within the same trial type, and a similar trial protocol was followed. Experimental tanks were cleaned, and water was changed before switching residency for each trial type and before starting the protocol for a new trial type.

The details of the trials are provided below and are explained in Figure 1.

**Figure 1.**
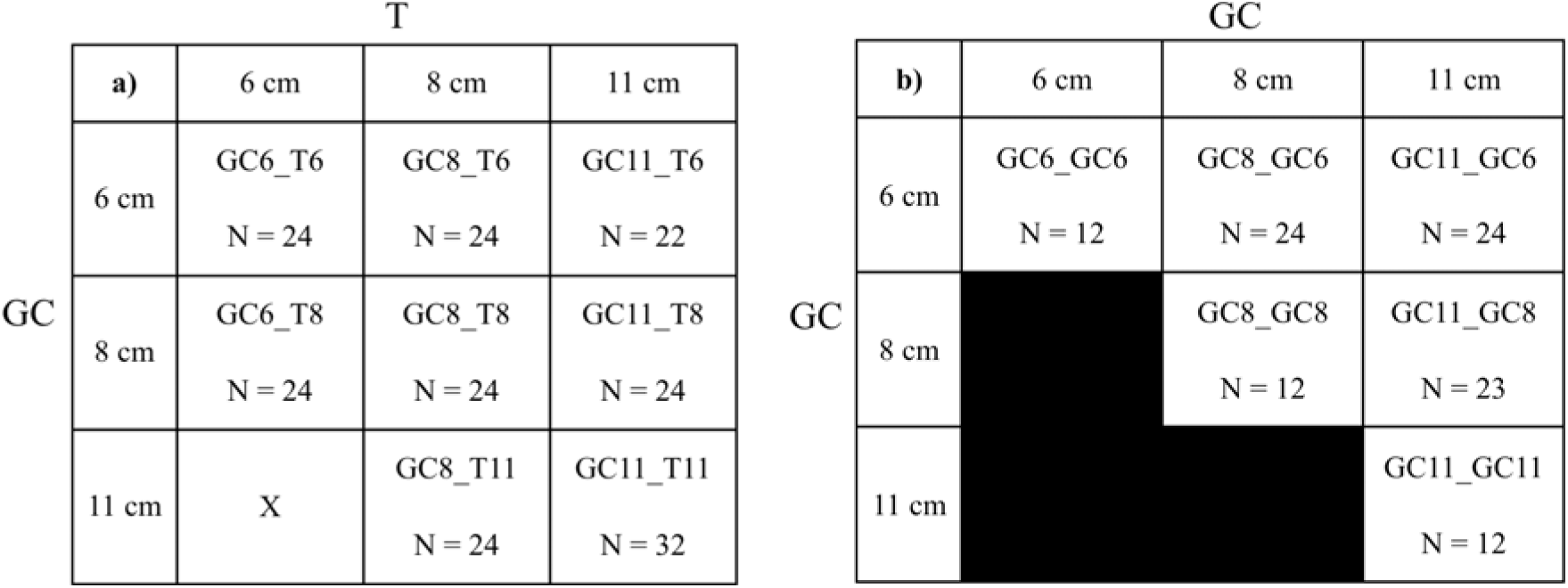
Schematic representation of a) interspecific and b) intraspecific trials. GC refers to green chromide, while T refers to tilapia. 6 cm, 8 cm, and 11 cm indicate the size classes (total length in cm). Each cell represents a given trial type. Sample size (N) represents the total number of trials (including both pre- and post-switching, R and I roles) for a particular trial type. Role reversal was not applicable for the size-matched intraspecific trials.

#### 2.2.1 Interspecific Trials

Interspecific trials involved pairing one GC with one T. GC11_T11, GC8_T8, and GC6_T6 represented size-matched trials (Figure 1). Size-unmatched trials consisted of pairings between different size classes, namely GC11_T6, GC11_T8, GC8_T11, GC8_T6, and GC6_T8 (Figure 1). We could not conduct trials for GC6_T11 due to the high aggression shown by T11 towards GC6, which resulted in the latter being distressed.

Each of the size-matched and size-unmatched trials, except GC11_T11 and GC11_T6, had three biological replicates (3 pairs of GC and T in separate experimental tanks), and each biological replicate had 8 technical replicates (the same pair of fish repeated 8 times, where in, four trials were conducted over four consecutive days, providing a rest period of 24 h in between the trials. After this, the remaining four were conducted in a similar way but after switching the R and I roles for the given pair and giving a rest period of 48 h before swapping the roles). This gave rise to N=24 trials for a given trial type. An exception to this was GC11_T11, which had four biological replicates, and GC11_T6, where we lost two trials owing to a technical glitch in the recorder. In total, we had a sample size of 198 trials (GC11_T11: 32, GC8_T8: 24, GC6_T6: 24, GC8_T11: 24, GC11_T8: 24, GC11_T6: 22, GC8_T6: 24, GC6_T8: 24) (Figure 1).

#### 2.2.2 Intraspecific trials

Intraspecific trials were performed for all three size classes of GC, including GC6, GC8, and GC11. In each trial, two GC individuals were paired, and similar to the interspecific trials, intraspecific trials were also divided into size-matched trials, for example, GC11_GC11, GC8_GC8, and GC6_GC6, and size-unmatched trial types, for example, GC8_GC6, GC11_GC6, and GC11_GC8. Like the interspecific trials, residency was switched for the size-unmatched trials following the same protocol as mentioned above, and thus having a sample size of N=24 for each trial type in the intraspecific category. An exception to this was GC11_GC8, where we had 23 instead of 24 trials due to a technical glitch during the recording. However, switching was not applicable for the size-matched ones, thus having a sample size of N=12 for each trial type (Figure 1).

The protocol for biological and technical replicates in the intraspecific trials was also similar to those mentioned for the interspecific trial category. In total, the sample size for intraspecific trials was 107 (GC6_GC6: 12, GC8_GC8: 12, GC11_GC11: 12, GC8_GC6: 24, GC11_GC6: 24, GC11_GC8: 23) (Figure 1).

### 2.3 Behavioural readouts and definition

We quantified behavioural readouts for each individual for a given pair, and all behaviours were scored from the playback of video recordings. A total of 9150-min of video recordings, across inter- and intraspecific trials, were observed and analysed. The different behaviours that we scored are: a) bite, an event behaviour in which the actor physically contacted the receiver with an open mouth (Supplementary video 1, Section 1 of Supplementary information 1); b) charge (Supplementary video 2, Section 1 of Supplementary information 1), an event behaviour in which the actor moved towards the receiver with a speed of 1.6±0.25 (Mean ± SEM) TL/second for T6, 1.8±0.22 TL/second for T8, 1.8±0.2 TL/second for T11, 1.6±0.19 TL/second for GC6, 1.5±0.2 TL/second for GC8, and 1.6±0.2 TL/second for G11. and c) chase (Supplementary video 3, Section 1 of Supplementary information 1), a state behaviour in which the actor followed the receiver, with a speed of 2.93±0.29 (Mean ± SEM) TL/ second for T8, 2.84 ±0.29 for T11, 1.5±0.2 for GC6, 1.7.5±0.24 for GC8, and 1.7±0.3 for GC11. T6 did not initiate chase behaviour in any of the trials. The speed of each of these behaviours was calculated by using the 8×8 cm grid (speed = number of grids covered by the fish/duration of the behaviour). For both charge and chase, speed was analysed using a subset of total incidents, pooled across R and I status, separately for GC6, GC8, GC11, T6, T8, and T11 across matched and unmatched trial types for both interspecific and intraspecific categories. The sample size for charge was 15 for all of these categories. However, for chase, the sample size was 15 for T8, T11, and GC8, 14 for GC11, and 12 for GC6. Bite and charge were event behaviours, and chase was state behaviour. However, we summed the incidences (number of times the behaviours, chase, bite, or charge occurred) of bite, chase, and charge and expressed them as total aggression for an individual in a given trial type. The next readout that we measured was feeding, an event behaviour, which was calculated as the number of feeding bouts. A feeding bout was defined when the actor moved towards the food particle, opened its mouth, and took a bite of the food (Supplementary video 4, Section 1 of Supplementary information 1). We also calculated the hiding duration. Hiding (Supplementary video 5, Section 1 of Supplementary information 1) was defined as a state behaviour in which the actor moved behind the filter and remained there for>1 second, with >50% of its body behind the filter.

### 2.4 Statistical analysis

#### 2.4.1 Factors influencing the outcome of interspecific contests - scoring

To quantify the outcomes of interspecific contests, we scored aggression, feeding, and hiding on a binary scale. Here, the individual who initiated the first aggressive act during the 30-min observation period got a score of 1. Similarly, the individual who fed first during the observation period also got a score of 1. However, the individual who hid first got a score of - 1. In all three cases, the opponent who did not initiate the behaviour received a score of 0. We then summed the scores for each individual, and the final score ranged from -1, 0, 1, up to a maximum of 2. Based on the scores, winners were decided, and scores of the winners were considered for further analysis. Trials in which two individuals tied were not considered. Similar scoring systems have also been developed in other studies, for example, to study intraspecific contests in the forked fungus beetle (*Bolitotherus cornutus)* (Mitchem et al., 2019) and in the ruin lizards *(Podarcis siculus)* (Titone et al., 2017), and for investigating interspecific competition between pearl cichlids (*Geophagus brasiliensis*) and Nile tilapia (*Oreochromis niloticus*) (Sanches et al., 2012). In our study, since the response variable, score of winners, was ordinal in nature, a cumulative link mixed model (CLMM) with a logit link function was used (Costa et al., 2018; Grønberg, 2023). The CLMM was fitted with the Laplace approximation to account for the random effects (the same pair was tested across 8 technical replicates). In the model, the fixed effects were species (GC/T), residency status (R/I), developmental stage (LL/MM/SS), and body size contrast (N/L/M/S). The developmental stage LL was assigned to score data (for both GC and T) from all the trials having 11cm individuals; similarly, MM included 8 cm individuals (both GC and T), and SS comprised 6 cm individuals (both GC and T), from all trial types. The body size contrast N (no contrast) was assigned to data (for both GC and T) across all size-matched trials, including GC11_T11, GC8_T8, and GC6_T6 trial types, wherein there was no contrast in body sizes. For the size-unmatched trial types, we divided the trials into three contrast categories, L, M and S. The contrast category, L (largest contrast), included GC11_T6 wherein the contestants were 6 and 11 cm, M (medium contrast) included GC11_T8 and GC8_T11 trial types wherein 8 and 11 cm individuals were paired, and S (small contrast) included GC8_T6 and GC6_T8 trial types wherein 8 and 6 cm individuals were paired. TankID was included as a random intercept to account for technical replicates. Initially, we had included an interaction between developmental stages and species, and residency and species; however, based on Akaike’s Information Criterion (AIC), the interaction term was removed, and the additive model was retained as it provided the best fit (AIC = 219.85). The reference levels used in the model were GC for species, LL for developmental stage, R for residency status, and L for body size contrast. Estimated marginal means were calculated using the emmeans package, and post hoc pairwise contrasts were performed for developmental stage and body size contrast, as they had more than two levels for each of the variables. All inferential analyses were conducted in R (version 4.5.0) using the packages readxl to read the data files, ordinal for the CLMM model, dplyr for data formatting, emmeans for post hoc comparisons, and ggplot2 for data visualization.

#### 2.4.2 Factors influencing the outcome of interspecific contests - absolute behaviours

Apart from scoring, we also analysed the absolute behavioural data on total aggression, feeding bouts, and hiding duration. Normality of the raw behavioural data was assessed using the Shapiro-Wilk tests, which indicated significant deviations from normality (P < 0.05) for all three behaviours. Thus, we used separate generalized linear mixed models (GLMMs) to analyse each of the three behaviours. However, the fixed effect remained the same as in the scoring analysis, namely, species (GC/T), residency status (R/I), developmental stage (LL/MM/SS), and body size contrast (N/L/M/S).

Total aggression and feeding bouts were analysed using a zero-inflated Generalised Linear Mixed Model (GLMM) with a negative binomial error distribution and a log link function to account for overdispersed count data and a high frequency of zero values (Mollier et al., 2017). For total aggression and feeding bouts’ models, in addition to the fixed effects as mentioned above, we included two interaction effects, species*developmental stage and species*residency. This was done to examine whether the effects of body size (or developmental stage) and residency status differed between the species (GC, T). We then chose the best model using the stepwise deletion approach, whereby non-significant interaction terms were sequentially removed, and competing models were compared using Akaike’s Information Criterion (AIC) (Crawley, 2013). The species*residency interaction was not significant and was therefore excluded from both the total aggression and feeding bouts’ model. In the final model, we retained the species*developmental stage interaction, together with the additive effects of residency status and body size contrast, and TankID was included as a random intercept to account for technical replicates (Total aggression/feeding bouts ∼ species*developmental stage + residency + body size contrast + (1 | tankID)). The selected models also had the lowest AIC (AIC = 2262.3 for total aggression and AIC = 3685.03 for the feeding bouts model) and therefore provided the best fit.

Hiding duration was analysed using a two-step hurdle approach (Álvarez-Ruiz et al., 2021; Ocepek et al., 2017) because it was a continuous variable, unlike total aggression and feeding bouts. It also contained a large number of zero values corresponding to trials in which individuals did not exhibit the hiding behaviour. Although a gamma distribution is commonly used to model positive continuous duration data, it can only accommodate values >0 (Chen et al., 2022). Thus, first, the occurrence of hiding (presence/absence) was analysed using a binomial GLMM with a logit link function. Second, for observations in which hiding occurred (Hiding>0 s), duration was analysed using a GLMM with a Gamma error distribution and a log link function. As with the total aggression and feeding models, for both components of the hurdle analysis, fixed effects initially included both the interaction terms, species*developmental stage and species*residency status, in addition to the main effects. We then simplified the models using the stepwise deletion approach, as mentioned above. The species*residency interaction was not significant for both components of the model and was therefore excluded. Similar to the other two models, the hiding duration model was defined as hiding duration ∼ species*developmental stage + residency + body size contrast + (1 | tankID), which also provided the best fit (AIC = 424.82 for the binomial model, AIC = 3157.7 for the gamma model).

For all three behavioural models, estimated marginal means were calculated using the emmeans package, and post hoc pairwise contrasts were performed. These contrasts were conducted on the model scale and back-transformed to the response scale, and the results are reported based on this. For the hiding duration model, post hoc comparisons are reported only for the second part of the GLMM model. Additionally, in all three models, the reference category was GC as I and LL in the developmental stage, and L type in the body size contrast, and all model effects are reported as deviations from these given reference categories. All the analyses were conducted in R (version 4.5.0) using the packages readxl to read the data files, glmmTMB for all the GLMM models, emmeans for post hoc comparisons, and ggplot2 and ggbreak for data visualization.

#### 2.4.3 Factors influencing the outcome of intraspecific contests - absolute behaviours

To understand how green chromide interacts with its conspecifics, we also analysed the absolute behavioural data on total aggression, feeding bouts, and hiding duration of the intraspecific trials. Normality of the raw behavioural data was assessed using the Shapiro-Wilk tests, which indicated significant deviations from normality (P<0.05) for all three behaviours. Thus, we used separate generalized linear mixed models (GLMMs) to analyse each of the three behaviours. In this case, the fixed effects, except species, remained the same as in the interspecific analyses, thus including residency status (R/I), developmental stage (LL/MM/SS), and body size contrast (N/L/M/S).

Total aggression and feeding bouts were analysed using a zero-inflated Generalised Linear Mixed Model (GLMM) with a negative binomial error distribution and a log link function to account for overdispersed count data and high frequency of zero values. An additive model was used for both behaviours, wherein, along with the fixed effects, TankID was included as a random intercept to account for technical replicates (Total aggression/feeding bouts ∼ developmental stage + residency + body size contrast + (1 | tankID)). Both models provided a good fit, with an AIC of 1658.7 for the total aggression model and 1990.4 for the feeding bouts model.

Just like the hiding duration data for the interspecific model, the intraspecific trials also contained several zero values. Thus, we used a two-step hurdle approach for this case as well. Similar to the interspecific model, the occurrence of hiding (presence/absence) was analysed using a binomial GLMM with a logit link function, and then, for observations in which hiding occurred (Hiding>0 s), duration was analysed using a GLMM with a Gamma error distribution and a log link function. As with the total aggression and feeding models, for both components of the hurdle analysis, we used an additive model, which included TankID as a random intercept. The model was defined as hiding ∼ developmental stage + residency + body size contrast + (1 | tankID), and it provided a good fit (AIC = 140.7 for the binomial model, AIC = 1698.2 for the gamma model).

In all three models, the reference category was R in the residency status, LL in the developmental stage, and L in the body size contrast, and all model effects are reported as deviations from these given reference categories. All the analyses were conducted in R (version 4.5.0) using the packages readxl to read the data files and glmmTMB for all the GLMM models.

#### 2.4.4 Comparing GC in interspecific versus intraspecific trials

To assess the performance of GC in inter-versus intraspecific trials, we kept the size class or developmental stage of GC constant across such comparisons. For example, GC11 was assigned to all 11 cm individuals of GC belonging to GC11_T11, GC11_T8, and GC11_T6 interspecific trials and was compared to the performance of GC11 individuals in intraspecific trials, including GC11_GC11, GC11_GC8, and GC11_GC6 trial types. Similarly, the performance of GC8 individuals in interspecific trials included data from GC8_T11, GC8_T8, and GC8_T6 trials and was compared with the GC8 individuals in intraspecific trials, including data from GC8_GC8, GC11_GC8, and GC8_GC6 trials. Finally, behavioural data for GC6 individuals included GC6_T8 and GC6_T6 interspecific trials and were compared with the performance of GC6 individuals in intraspecific trials, including GC6_GC6, GC11_GC6, and GC8_GC6 trials. The comparisons were done separately for total aggression, feeding, and hiding data. Since the behavioural data from both interspecific and intraspecific trials significantly deviated from normality, comparing the performance of GC in inter-versus intraspecific trials was conducted using the pairwise (two-tailed) Wilcoxon rank-sum tests. All statistical tests were conducted in R (version 4.5.0), and the significance was assessed at α = 0.05.

### 2.5 Ethical note

All procedures conducted in this study were reviewed and approved by the University Research Board and the Institute Biosafety Committee under project code AU/SUG/SAS/BLS/2018–19/20. Although we sought for permits, no field collection permits were required as all the authors are Indian nationals conducting purely academic, non-commercial laboratory research, which is exempt from prior approval under Section 7 of The Biological Diversity Act (2002) (Ministry of Law and Justice, 2003). Furthermore, because both species are widely designated as food fish and aquaculture species rather than protected wildlife under the Wild Life (Protection) Act, 1972, no specific wildlife propagation permits were necessary to use them in laboratory behavioural studies (Government of India, 2023).

The experiments in this study were only observational and no invasive procedures were performed. Additionally, all behavioural trials were conducted under controlled laboratory conditions adhering to the national statutory framework established by the CPCSEA Guidelines for Experimentation on Fishes [Committee for the Purpose of Control and Supervision of Experiments on Animals (CPCSEA), 2021] and the international guidelines (Association for the Study of Animal Behaviour Ethical Committee & Animal Behavior Society Animal Care Committee [ASAB/ABS], 2025). Fish were monitored daily to assess overall health, and daily checks were done to ensure that the filtration systems were properly functioning. Fish were transferred between home and experimental tanks gently using hand-held nets. To minimize handling stress, all procedures were conducted under low light conditions.

Aggressive interactions observed during the trials consisted primarily of bites, chases, and charges, and were generally brief. No physical injuries were observed in the fish during or following the majority of trials. In order to protect the welfare of the fish, we decided that trials would be terminated if aggression became excessive or prolonged, particularly if repeated attacks posed a risk of physical injury, severe distress, or the receiver of aggression being cornered. During one set of five trials, GC6_T11 (wherein the oldest developmental stage of T was paired with the youngest developmental stage of GC), aggression was considered excessive and the trials were therefore aborted. As a precautionary measure, this entire trial type was subsequently discontinued. No mortality occurred during the study, and all the fish were returned to their home tanks in good condition, once the experiments were completed and are still maintained in the laboratory to date.

## 2. Results

### 3.1 Factors influencing the outcome of interspecific contests - Scoring

Of the 198 trials, 17 ended in a tie, leaving 181 trials with a clear winner, and the winners either had a score of 1 or 2. Out of the 181 trials, GC won in 39 trials, whereas T won in 142 trials. Similarly, R won 110 contests, while I won only 71 contests. Across developmental stages, LL, MM, and SS won 68, 65, and 48 trials, respectively. The distribution of winners also varied across body size contrast categories, with L, M, S, and N having 20, 45, 45, and 71 winners or trials, respectively.

The cumulative link mixed model, wherein score was the response variable and species, developmental stage, residency status, and body size contrast were the fixed effects, converged successfully.

#### 3.1.1 The species effect

Species had a significant effect on the scores, wherein among the winners, T was significantly more likely to have higher scores compared to GC (β=2.17 ± 0.62, P=0.000475) (Table 1).

**Table 1.**
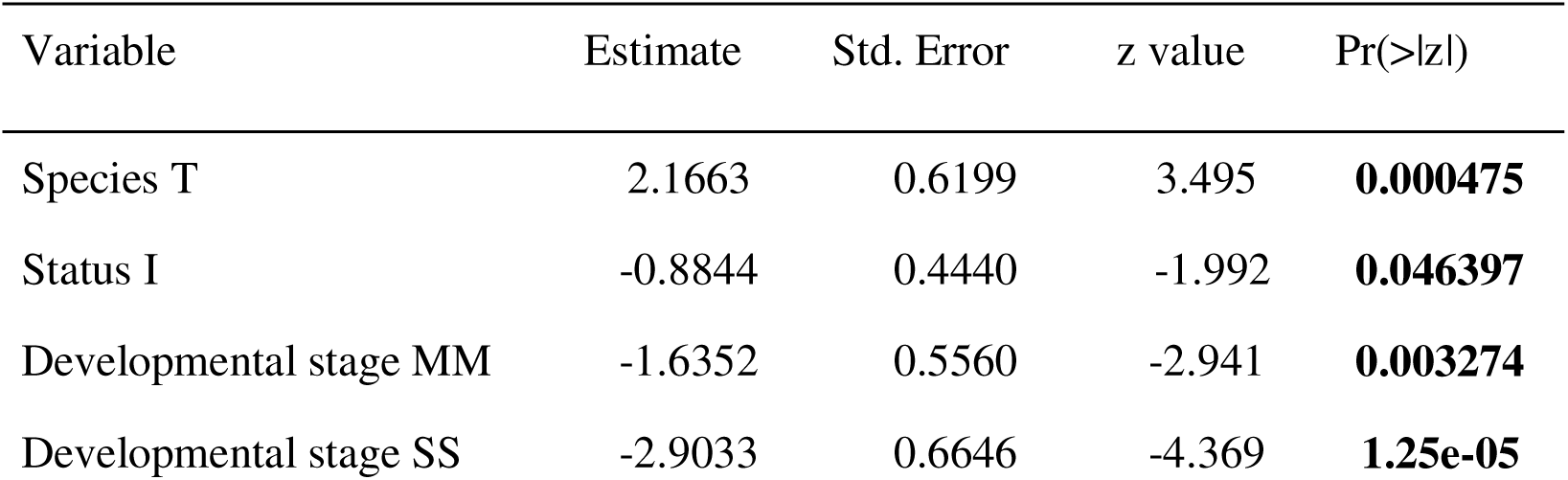
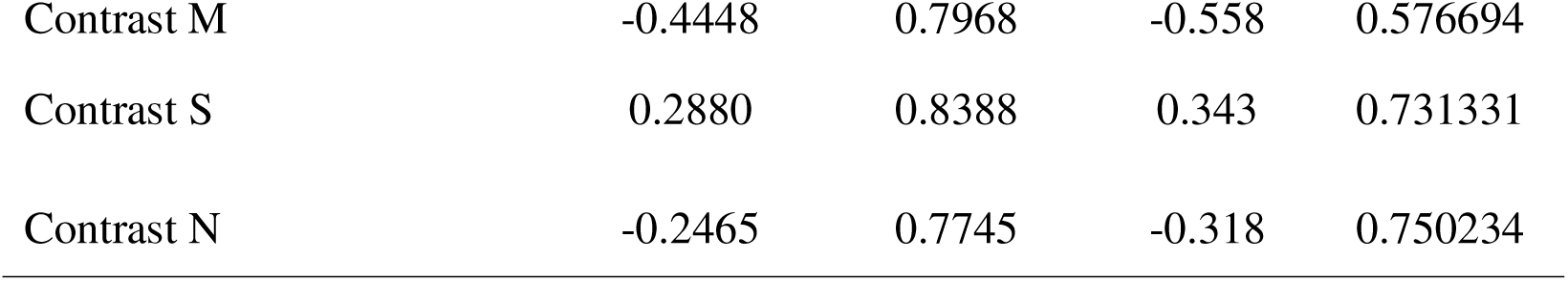
Summary of the CLMM model for the scoring data. P<0.05 is indicated in bold.

#### 3.1.2 The residency effect

Residency status also had a significant effect on the scores, with I winners significantly less likely to have higher scores than R (β=-0.88 ± 0.44, P=0.046397) (Table 1).

#### 3.1.3 The effect of developmental stages

The developmental stage had a significant effect on the scores. Both MM (β=-1.63 ± 0.55, P=0.003274) and SS winners (β=-2.90 ± 0.66, P=1.25e-05) were significantly different when compared to LL winners (Table 1). Post hoc pairwise comparisons further revealed that while MM (P=0.0092) and SS (P<0.0001) had significantly lower scores than LL, there was no significant difference between MM and SS (P=0.5385).

#### 3.1.4 The effect of body size contrast

None of the body size contrasts had a significant effect on the scores. The winners from M (β=-0.44 ± 0.79, P=0.576694), S (β=0.28 ± 0.83, P=0.731331), and N (β=-0.24 ± 0.77, P=0.750234) did not significantly differ when compared to L (Table 1).

### 3.2 Factors influencing the outcome of interspecific contests - Absolute behaviours

Our behavioural data showed that T11, as R, had the highest value of total aggression (Mean = 135, Range = 7-274) in GC8_T11 trials (Table S1 in Section 2 of the supplementary information 1). However, for feeding bouts, the highest values were recorded for GC, as I, in GC8_T8 (Mean = 184, Range = 48-415) trials (Table S1 in Section 2 of the supplementary information 1). However, GC also had zero values of feeding in several trial types, including GC6_T6, GC8_T6, GC6_T8, GC8_T11, and GC11_T11, while T had zero values in two of the trial types, including GC8_T6 and GC8_T11 (Table S1 in Section 2 of the supplementary information 1). The highest value for hiding duration was for GC as I in GC8_T11 (Mean = 929.41, Range = 23-1819); contrastingly, much higher than T, which had the highest value of hiding as R in GC11_T6 (Mean = 90.64, Range = 0-670) (Table S1 in Section 2 of the supplementary information 1). The complete mean and range values for GC and T as R and I from all trial types are provided in Table S1 in Section 2 of the Supplementary Information 1.

The results of the GLMM models for all three behaviours are given below.

#### 3.2.1 Total aggression

In the total aggression model, species and residency status showed a significant main effect, wherein T (β=2.812 ± 0.386, P=3e-13) was significantly more aggressive than GC, and R (β=0.626 ± 0.126, P=6e-07) showed higher aggression than I (Table 2, Figure 2). On the other hand, none of the developmental stages (MM: β=-0.463 ± 0.263, P=0.079; SS: β-0.564 ± 0.305, P=0.064) and body size contrasts (M: β = -0.785 ± 0.489, P=0.108; N: β=-0.203 ± 0.448, P=0.651; S: β=-0.286 ± 0.484, P>0.555) had a significant effect on total aggression (Table 2, Figure 2). Interestingly, the interaction terms had a significant effect on the data, wherein both SpeciesT*DevelopmentalStageMM (β=-3.485±0.47, P=1e-13) and SpeciesT*DevelopmentalStageSS (β = -3.964±0.486, P=3e-16) showed significantly lower aggression than the reference category (SpeciesGC*DevelopmentalStageLL) (Table 2).

**Figure 2.**
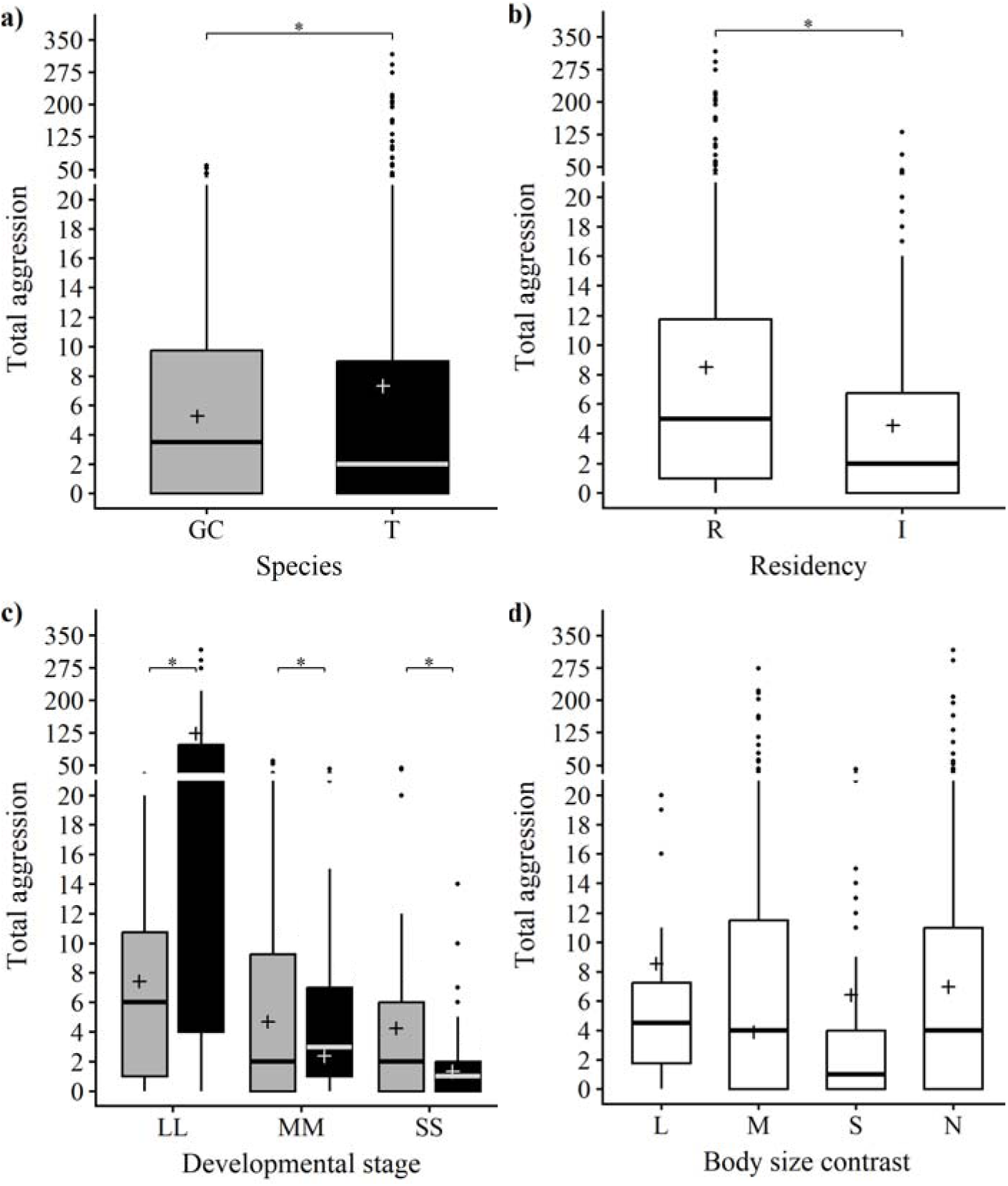
Behavioural data on total aggression as per the a) species identity, where 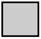 represents green chromides (GC) and 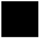 represents Tilapia (T); b) residency, where R is resident, and I is intruder; c) developmental stages where LL represents 11 cm individuals, MM represents 8 cm individuals, and SS represents 6 cm individuals, and 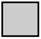 represents green chromides (GC) and 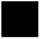 represents Tilapia (T); and d) body size contrasts, where L represents trial types with the largest contrast (11 cm and 6 cm individuals paired), M represents medium contrast (11 cm and 8 cm individuals paired), S represents small contrast (8 cm and 6 cm individuals paired), and N represents no contrast (size-matched individuals paired). The line inside each box represents the median, the box denotes the interquartile range, the whiskers indicate the minimum and the maximum values, and dots indicate outliers. ‘+’ represents the model estimated marginal mean values backtransformed to the response scale, and ‘*’ indicates significant (P<0.05) differences between boxes connected by shoulder bars (post-hoc comparisons of GLMM model).

**Table 2.** Estimates of interspecific GLMM models for aggression, feeding, and hiding behaviours.

| Variable | Estimate | Std. Error | z value | Pr(>\|z\|) |
| --- | --- | --- | --- | --- |
| **Aggression: Negative binomial GLMM** |  |  |  |  |
| Intercept | 2.009 | 0.386 | 5.2 | **2e-07** |
| Species T | 2.812 | 0.384 | 7.318 | **3e-13** |
| Developmental stage MM | -0.463 | 0.263 | -1.76 | 0.079 |
| Developmental stage SS | -0.564 | 0.305 | -1.85 | 0.064 |
| Residency Status R | 0.626 | 0.126 | 4.976 | **6e-07** |
| Contrast M | -0.785 | 0.489 | -1.61 | 0.108 |
| Contrast N | -0.203 | 0.448 | -0.45 | 0.651 |
| Contrast S | -0.286 | 0.484 | -0.59 | 0.555 |
| Species T * Developmental Stage MM | -3.485 | 0.47 | -7.41 | **1e-13** |
| Species T * Developmental Stage SS | -3.964 | 0.486 | -8.16 | **3e-16** |
| **Feeding: Negative binomial GLMM** |  |  |  |  |
| Intercept | 4.07 | 0.191 | 21.26 | **2e-100** |
| Species T | -0.423 | 0.166 | -2.551 | **0.011** |
| Developmental stage MM | -0.345 | 0.157 | -2.202 | **0.028** |
| Developmental stage SS | -1.416 | 0.169 | -8.386 | **5e-17** |
| Residency Status R | 0.185 | 0.07 | 2.636 | **0.008** |
| Contrast M | 0.013 | 0.236 | 0.054 | 0.957 |
| Contrast N | 0.282 | 0.221 | 1.277 | 0.201 |
| Contrast S | 0.256 | 0.239 | 1.073 | 0.283 |
| Species T * Developmental Stage MM | 0.218 | 0.22 | 0.989 | 0.323 |
| Species T * Developmental Stage SS | 1.235 | 0.224 | 5.509 | **4e-08** |
| **Hiding occurrence: Binomial GLMM** |  |  |  |  |
| Intercept | -0.31 | 0.747 | -0.414 | 0.679 |
| Species T | -1.786 | 0.578 | -3.089 | **0.002** |
| Developmental stage MM | 1.531 | 0.562 | 2.725 | **0.006** |
| Developmental stage SS | 6.179 | 1.319 | 4.686 | **3e-06** |
| Residency Status R | 0.098 | 0.292 | 0.336 | 0.737 |
| Contrast M | 0.883 | 0.915 | 0.966 | 0.334 |
| Contrast N | -0.525 | 0.82 | -0.64 | 0.522 |
| Contrast S | -0.619 | 0.929 | -0.666 | 0.505 |
| Species T * Developmental Stage MM | 2.149 | 0.835 | 2.574 | **0.01** |
| Species T * Developmental Stage SS | -2.723 | 1.317 | -2.067 | **0.039** |
| **Hiding duration: Gamma GLMM** |  |  |  |  |
| Intercept | 5.678 | 0.57 | 9.969 | **2e-23** |
| Species T | -3.44 | 0.447 | -7.69 | **1E-14** |
| Developmental stage MM | 0.173 | 0.362 | 0.478 | 0.632 |
| Developmental stage SS | 1.948 | 0.451 | 4.318 | **2e-05** |
| Residency Status R | 0.306 | 0.183 | 1.678 | 0.093 |
| Contrast M | 0.425 | 0.511 | 0.831 | 0.406 |
| Contrast N | -0.347 | 0.464 | -0.75 | 0.455 |
| Contrast S | -0.671 | 0.434 | -1.55 | 0.122 |
| Species T * Developmental Stage MM | 1.763 | 0.579 | 3.045 | **0.002** |
| Species T * Developmental Stage SS | 0.184 | 0.565 | 0.326 | 0.745 |

517 P < 0.05 is indicated in bold

Post hoc analysis revealed that while GC was significantly more aggressive than T in the MM (P=0.00196) and SS (P=2.9e-05) developmental stages, T was significantly more aggressive than GC in the LL stage (P=2.5e-13) (Figure 2).

#### 3.2.2 Feeding bouts

In the feeding model, species and residency status showed a significant main effect, wherein T showed significantly lower (β=-0.423 ± 0.166, P=0.011) and R (β=0.185 ± 0.07, P=0.008) showed significantly higher feeding than GC and I, respectively (Table 2, Figure 3). Additionally, both MM (β=-0.345 ± 0.157, P=0.028) and SS (β=-1.416 ± 0.169, P=5e-17) developmental stages fed significantly less than LL. On the other hand, none of the body size contrasts (M: β=0.013 ± 0.236, P=0.957; N: β=0.282 ± 0.221, P=0.201; S: β=0.256 ± 0.239, P=0.283) had a significant effect on the feeding behaviour (Table 2, Figure 3). The interaction term SpeciesT*DevelopmentalStageMM (β=0.218 ± 0.22, P=0.323) did not have a significant effect on the response variable; however, SpeciesT*DevelopmentalStageSS (β=1.235 ± 0.224, P=4e-08) showed significantly higher feeding than the reference category (SpeciesGC*DevelopmentalStageLL) (Table 2).

**Figure 3.**
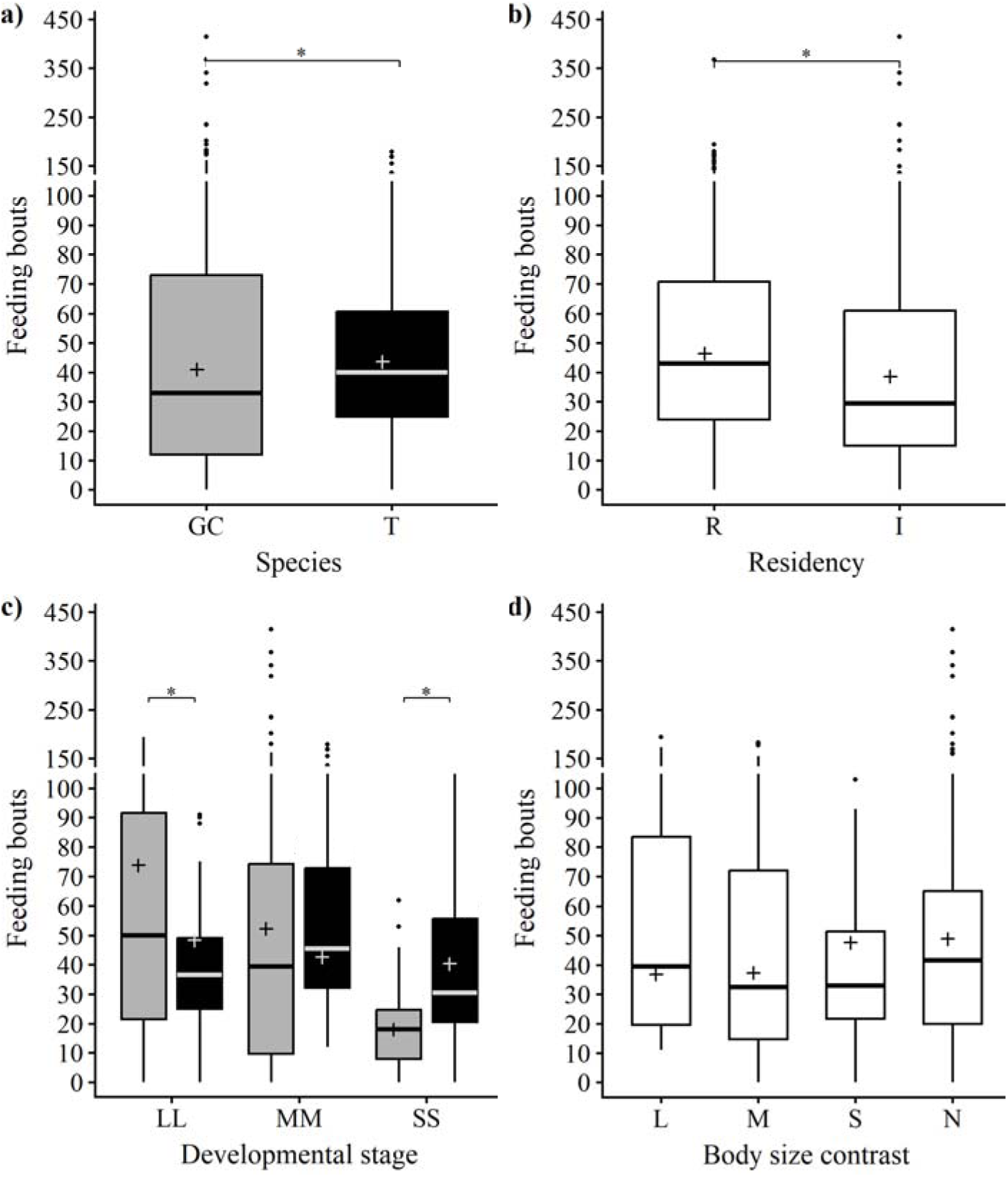
Behavioural data on feeding bouts as per the a) species identity, where 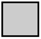 represents green chromides (GC) and 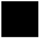 represents Tilapia (T); b) residency, where R is resident, and I is intruder; c) developmental stages, where LL represents 11 cm individuals, MM represents 8 cm individuals, and SS represents 6 cm individuals, and 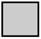 represents green chromides (GC) and 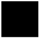 represents Tilapia (T); and d) body size contrasts, where L represents the trial types with the largest contrast (11 cm and 6 cm individuals paired), M represents medium contrast (11 cm and 8 cm individuals paired), S represents small contrast (8 cm and 6 cm individuals paired), and N represents no contrast (size-matched individuals paired). The line inside each box represents the median, the box denotes the interquartile range, the whiskers indicate the minimum and the maximum values, and dots indicate outliers. ‘+’ represents the model estimated marginal mean values backtransformed to the response scale , and ‘*’ indicates significant (P<0.05) differences between boxes connected by shoulder bars (post-hoc comparisons of GLMM model).

Post hoc comparisons revealed that, while GC fed significantly more than T in the LL (P=0.011) category, T fed significantly more than GC in the SS category (P=4e-08). There was no significant difference between GC and T in the MM category (P=0.114) (Figure 3).

#### 3.2.3 Hiding duration

In the case of hiding duration, the first part of the hurdle analysis, i.e, the binomial GLMM model, revealed that species had a significant effect on the response variable, wherein T was less likely to hide (T had significantly more zero values) than GC (β=-1.786 ± 0.578, P=0.002). However, residency (β=0.098 ± 0.292, P=0.737) and body size contrasts (M: β=0.883 ± 0.915, P=0.334; N: β=-0.525 ± 0.82, P=0.522; S: β=-0.619 ± 0.929, P=0.505) did not have a significant effect in the model (Table 2). Additionally, MM (β=1.531 ± 0.562, P=0.006) and SS (β=6.179 ± 1.319, P=3e-06) developmental stages had a significant effect in the model, wherein both were more likely to hide as compared to LL (LL had significantly more zero values for hiding durations) (Table 2). Finally, both the interaction terms had a significant effect, wherein SpeciesT*DevelopmentalStageMM (β=2.149±0.835, P=0.01) had significantly more non-zero values of hiding durations, whereas SpeciesT*DevelopmentalStageSS (β=–2.732 ± 1.317, P=0.039) had significantly fewer non-zero values (so higher zero values) when compared to the reference category (SpeciesGC*DevelopmentalStageLL) (Table 2).

The second part of the hurdle analysis, the Gamma GLMM model, revealed that species had a significant effect on hiding durations, wherein T hid significantly less than GC (β=-3.44 ± 0.447, P=1e-14); however, neither residency (β=0.306 ± 0.183, P=0.093) nor body size contrast (M: β=0.425 ± 0.511, P=0.406; N: β=-0.347 ± 0.464, P=0.455; S: β=-0.671 ± 0.434, P=0.122) had a significant effect in the model (Table 2, Figure 4). Additionally, the SS (β=1.948 ± 0.451, P=2e-05) developmental stage had a significantly longer hiding duration, while the MM (β=0.173 ± 0.362, P=0.632) developmental stage did not differ significantly from the LL category (Table 2, Figure 4). Lastly, while the SpeciesT*DevelopmentalStageMM had a significantly (β=1.763 ± 0.579, P=0.002) longer hiding duration, SpeciesT*DevelopmentalStageSS (β=0.184 ± 0.565, P=0.745) did not have a significant difference, both when compared to the reference category, SpeciesGC*DevelopmentalStage LL (Table 2).

**Figure 4.**
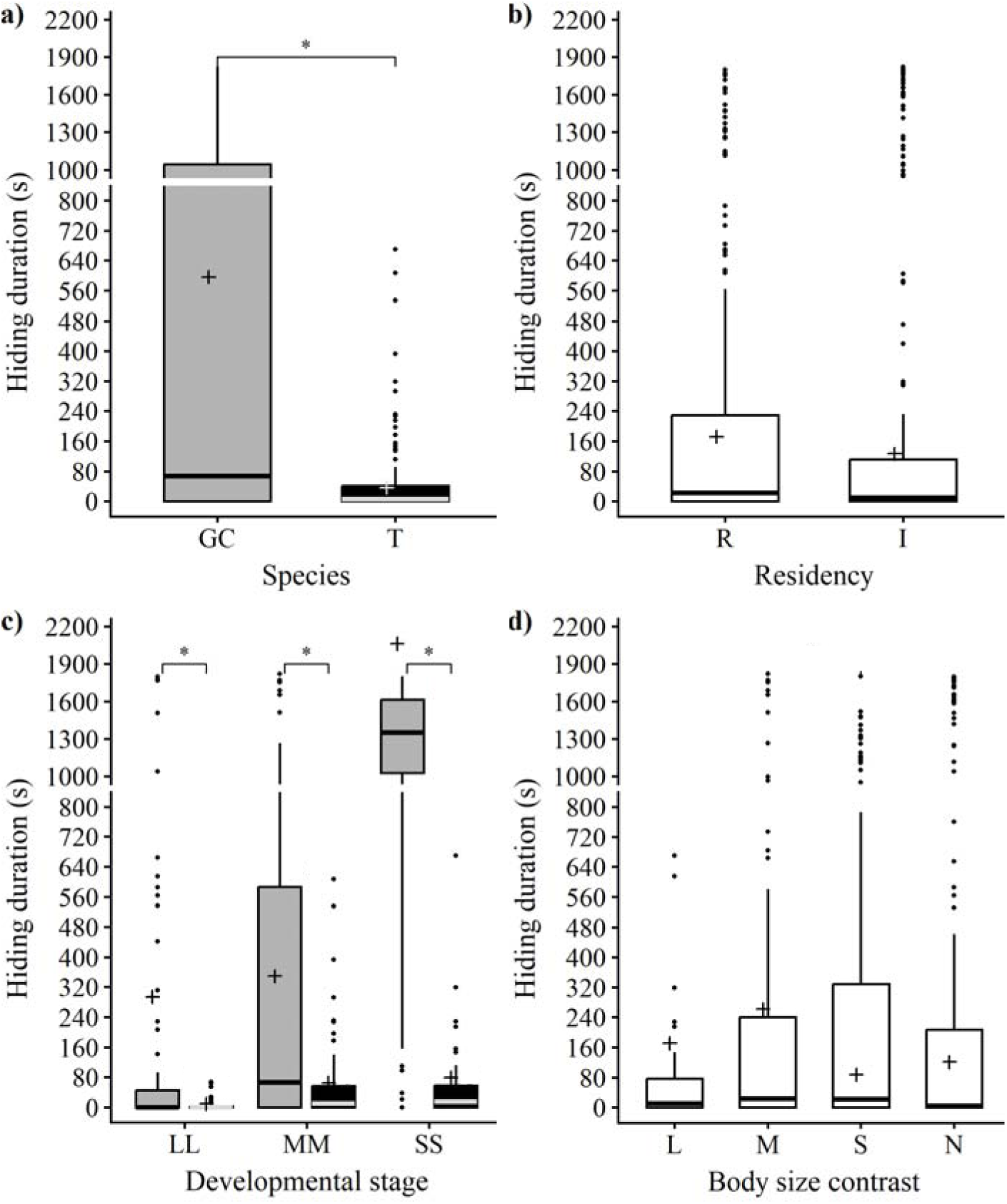
Behavioural data on hiding duration(s) as per the a) species identity, where 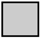 represents green chromides (GC) and 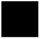 represents Tilapia (T); b) residency, where R is resident, and I is intruder; c) developmental stage, where LL represents 11 cm individuals, MM represents 8 cm individuals, and SS represents 6 cm individuals, and 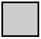 represents green chromides (GC) and 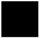 represents Tilapia (T); and d) body size contrast where L represents the trial types with the largest contrast (11 cm and 6 cm individuals paired), M represents medium contrast (11 cm and 8 cm individuals paired), S represents small contrast (8 cm and 6 cm individuals paired), and N represents no contrast (size-matched individuals paired). The line inside each box represents the median, the box denotes the interquartile range, the whiskers indicate the minimum and the maximum values, and dots indicate outliers. ‘+’ represents the model-estimated marginal mean values from the Gamma GLMM model backtransformed to the response scale, and ‘*’ indicates significant (P<0.05) differences between boxes connected by shoulder bars (post-hoc comparisons of the GLMM model).

Post hoc analysis showed that GC hid for a significantly longer duration than T in LL (P=1e-14), MM (P=1e-08), and SS (P=2e-20) categories (Figure 4).

### 3.3 Factors influencing the outcome of intraspecific contests - Absolute behaviours

In the case of intraspecific contests, our behavioural data showed that total aggression was highest for GC11 as R in GC11_GC11, with a value of 283 (Mean = 58.8, Range = 5-283). Similarly, feeding was also highest for GC11 as R in GC11_GC11 trials, having a value of 265 (Mean = 52.9, Range = 2-265). However, hiding duration was the highest for GC6 as R in GC6_GC6 trials, with a duration of 1702 seconds (Mean = 471.65, Range = 2-1702).

The results of the GLMM models for all three behaviours are given below, and the complete model output tables are given in Table S2, S3, and S4 in Section 2 of the supplementary information 1.

#### 3.3.1 Total aggression

In the total aggression model, residency status showed a significant main effect, wherein GC as I (β=-0.70522 ± 0.282799, P=0.012641) was significantly less aggressive than GC as R (Table S2 in Section 2 of the supplementary information 1). Additionally, while the SS (β=-1.0446 ± 0.348205, P=0.0027) developmental stage showed significantly lower aggression than LL, MM (β=-0.64879 ± 0.495886, P=0.190756) did not differ significantly from LL (Table S2 in Section 2 of the supplementary information 1). Finally, in terms of body size contrast, while N (β=0.806054 ± 0.404226, P=0.046145) showed significantly higher aggression than L, both M (β=-0.68523 ± .467483, P=0.142709) and S (β=0.399056 ± 0.528108, P=0.44987) did not have any significant effect on the response variable (Table S2 in Section 2 of the supplementary information 1).

#### 3.3.2 Feeding bouts

In the feeding bouts model, only the developmental stage showed a significant effect on the response variable, with both MM (β=-1.01819 ± 0.199692, P=3.42E-07) and SS (β=-0.97337 ± 0.164178, P=3.05E-09) feeding significantly less than LL (Table S3 in Section 2 of the supplementary information 1). Residency status (β=-0.02349 ± 0.10868, P=0.8289) and none of the body size contrasts (M: β=-0.48287 ± 0.270549, P=0.074299, S: β=0.172215 ± 0.266973, P=0.518884, N: β=-0.234341 ± 0.10868, P=0.36046) had any significant effect on feeding bouts (Table S3 in Section 2 of the supplementary information 1).

#### 3.3.3 Hiding duration

In the case of hiding durations, the first part of the hurdle analysis, i.e, the binomial GLMM model, revealed that developmental stage had a significant effect on the response variable, wherein both MM (β=8.460426 ± 1.82463, P=3.54E-06) and SS (β=4.325667 ± 0.960802, P=6.73E-06) were more likely to hide (had significantly fewer zero values) than LL (Table S4 in Section 2 of the supplementary information 1). However, residency (β=-1.03852 ± 0.712106, P=0.144735) did not have a significant effect on the response variable. In terms of body size contrast, while M (β=-6.16503 ± 2.134772, P=0.003878) was significantly less likely to hide compared to L, S (β=-1.49098 ± 1.50806, P=0.322821), and N (β=-0.947 ± 1.434629, P=0.509187) did not have a significant effect on the occurrence of hiding (Table S4 in Section 2 of the supplementary information 1).

The second part of the hurdle analysis, the Gamma GLMM model, revealed that none of the metrics, which were developmental stage (MM: β=0.519636 ± 0.721585, P=0.471444, SS: β=-0.07014 ± 0.612205, P=0.908782), residency status (β=0.001232 ± 0.267281, P=0.996323), and body size contrast (M: β=0.153344 ± 0.901196, P=0.864887, S: β=-0.93012 ± 0.745164, P=0.211953, N: β=-0.14572 ± 0.691311, P=0.833051), had any significant effect on hiding duration (Table S4 in Section 2 of the supplementary information 1).

### 3.4 Comparing GC in interspecific versus intraspecific trials

For aggression, GC6 (W = 1134, P = 0.0013) and GC11 (W = 1562.5, P=4.28e-06) showed significantly lower aggression in interspecific trials when compared to intraspecific trials (Figure 5a). However, no significant difference in aggression was detected in GC8 between interspecific and intraspecific trials (W = 2149.5, P = 0.092) (Figure 5a). For feeding, GC6 fed significantly less in interspecific trials compared to intraspecific trials (W = 1028.5, P = 0.00018), whereas GC8 (W = 2999, P = 0.073) and GC11 (W = 2604, P = 0.531) did not show any significant differences (Figure 5b). Hiding duration was significantly longer for GC6 (W = 2983, P=1.72e-11) and GC11 (W = 3679.5, P=1.8e-05) in interspecific trials when compared to intraspecific trials, while the duration for GC8 in interspecific trials was not significantly different when compared to intraspecific trials (W = 2589, P = 0.895) (Figure 5c).

**Figure 5.**
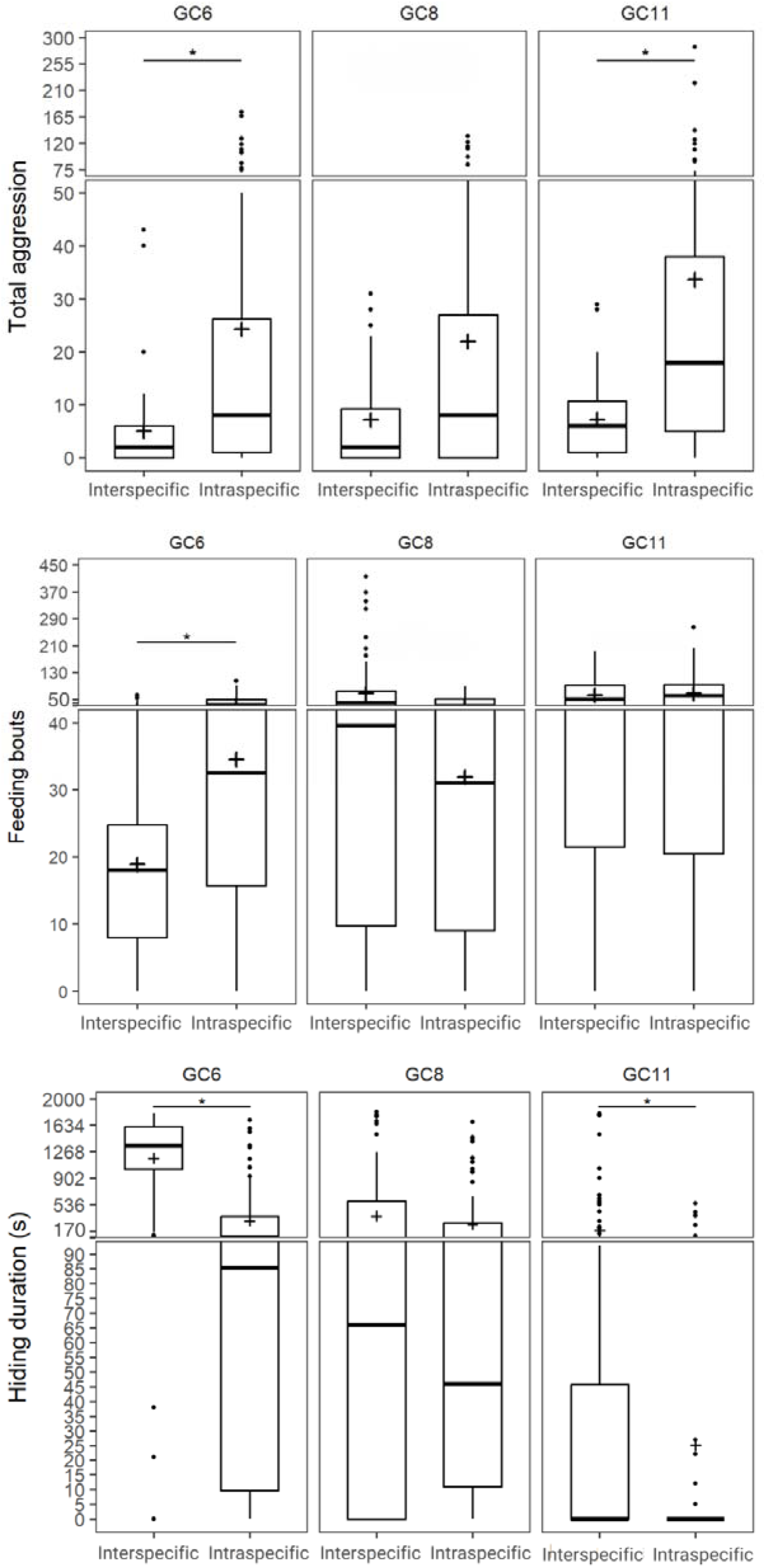
Behavioural performance of GC in interspecific versus intraspecific trials in terms of a) total aggression, b) feeding bouts, and c) hiding duration (s). The line inside each box represents the median, the box denotes the interquartile range, the whiskers indicate the minimum and the maximum values, and the dots denote the outliers. ‘+’ indicates the mean and ‘*’ indicates P<0.05 between boxes connected by shoulder bars, as indicated by the Wilcoxon rank-sum tests.

## 3. Discussion

In this study, we investigated how GC and T negotiated interference competition when facing each other. We primarily focused on GC and highlighted the context-dependent competitive strategies of the species. We showed that GC modulated its behavioural (aggression, feeding and hiding) responses, possibly to avoid fitness costs associated with direct physical confrontation, when encountering the aggressive T, as opposed to when interacting with conspecifics. Apart from the species effect, our results also suggest that residency status plays a significant role in governing competitive strategies (both aggression and feeding) for both the species, while developmental stage had a significant effect only on the feeding and hiding behaviours of the species. Interestingly, body size contrast, despite its established importance in previously documented literature (Arnott & Elwood, 2009; Chapin et al., 2019), played no significant role in behavioural dynamics. Our scoring data and absolute behavioural analysis revealed a very similar picture, wherein T emerged to be a winner in most of the interference trials, with a significantly higher level of total aggression. Our data also showed that GC is capable of exhibiting aggression; however, it strategizes and modifies the levels based on the characteristics of the opponent, for example, reduced aggression in inter-versus intraspecific trials. Interestingly, the feeding behaviour of GC was not substantially impacted in interspecific contests, and this could be because of the defence (hiding) strategy that GC exhibited to possibly avoid aggression in interspecific trials, while balancing hiding and feeding behaviours. Together, these findings suggest that GC is strategic in adjusting its fitness costs, emphasizing the adaptive traits of the species that may contribute towards its survival. However, there still remains a concern, due to the consistently high scores of T, along with its ability to show high levels of aggression at older developmental stages (T11), the large-scale invasion of T (Dwivedi & Mayank, 2021; Bera et al., 2023) may still threaten the long-term sustenance of GC.

One of the most unexpected findings of our study was that body size contrast between the opponents did not significantly influence the contest outcomes, and neither impacted the expression of competitive behaviours. Larger individuals are generally expected to possess a greater resource-holding potential (Arnott & Elwood, 2009; Martin & Ghalambor, 2014; Umbers et al., 2012), while contests between size-matched opponents are often predicted to escalate because competitors have similar fighting abilities (Draud, 2004; Lobregat et al., 2019; Pinto et al., 2019). Contrary to these expectations, body size contrast did not emerge as a significant predictor in either the scoring or behavioural analyses. This may be explained by three features of our system. First, aggression and more importantly, feeding remained prevalent even in size-unmatched contests, that too from the smaller-sized opponents, suggesting that asymmetric opponents were equally prepared to engage in competitive interactions, regardless of the size discrepancy. Second, the high values of total aggression displayed by T11 individuals, irrespective of the size of the opponents, likely masked any influence of body size asymmetry. Finally, hiding appeared to be largely species-specific, with GC consistently relying on this defensive behaviour more than T, suggesting that avoidance strategies were driven more by species identity than by relative body size. Consequently, the three factors: residency, species identity, and a marginal effect of developmental stage, explained competitive interactions more effectively than body size contrast, indicating that individual traits of the competitors, and not characteristics of the competition, had a significant role to play in governing the outcomes of the interspecific, interference competitions.

On the other hand, residency played a pivotal and consistent role in dictating competitive outcomes across all GC–T interaction types. Residents were more likely to win in terms of initiation of competitive behaviours, achieved higher scores, and showed more aggression and feeding in interspecific competition than intruders. In fact, resident GCs in the intraspecific trials also showed more aggression than intruders. The advantage of residency is generally attributed to the greater value that residents place on defending a space they already occupy, thus allowing them to exploit resources, for example, food, more efficiently (Kemp & Wiklund, 2004; O’Connor et al., 2015). Thus, residency may act as a metric of an opponent’s willingness to engage in competition and their ability to win. However, there was no significant effect of residency on hiding behaviour, and this may be because hiding was more strongly influenced by species identity, with GC significantly relying on this defensive behaviour in interspecific trials. Unlike residency, although developmental stage significantly affected the scores, its effect on total aggression was marginal and became significant only when species identity was considered. The ability of T11 to show higher levels of aggression, when compared to younger developmental stages of T and all developmental stages of GC, further suggests that aggressive responses may depend more on the specific identity (T species of a given developmental stage) of the opponent. In contrast, developmental stage significantly affected feeding regardless of species, consistent with previous studies showing that feeding can increase (Matsumura et al., 2021) or decrease (Lawson et al., 2018) with age depending on the species. The occurrence of zero feeding across several trial types, especially for GC (GC had zero values in four different trial types when compared to T, having zero values in two of the trial types), also suggests that interference competition can be a strong enough force to deter feeding altogether, potentially threatening the sustenance of the species. The developmental stage also affected hiding, although this pattern may be driven largely by GC, as T hardly hid regardless of the developmental stage. Interestingly, in intraspecific competition, developmental stage significantly affected all three behaviours, including aggression. This further strengthens the argument that unlike residency, developmental stage appears to be more context-dependent, with its effects varying with opponent identity.

Taken together, our findings suggest that the influence of assessment metrics on the expression of competitive behaviours depends on the competitor’s identity. This was particularly evident when we compared GC in inter- and intraspecific contests. Against conspecifics, GC generally invested more in aggression while relying less on hiding, when compared to competing with the highly aggressive T, suggesting that direct competition is favoured when opponents possess similar behavioural repertoires and competitive abilities (Peiman and Robinson., 2010). However, this behavioural shift did not really compromise resource acquisition. Despite encountering T11, which showed the highest aggression, GC maintained feeding, in both GC8 and GC11 stages. This suggests that GC does not simply withdraw from competition but instead reallocates its competitive efforts to maximize resource acquisition while minimizing the risks associated with direct conflict. However, it is important to note that we measured feeding bouts and not the amount of food consumed, which, if measured, may give a clearer idea of resource acquisition (Svanbäck & Bolnick, 2006; Rahman & Verdegem, 2010). One possible explanation of GC’s strategy in the presence of T is that hiding allows GC to avoid costly aggressive encounters while remaining close to the contested resource and exploiting feeding opportunities once aggressive interactions subside (Lanna et al., 2017; Hollis et al., 2004). Such behavioural flexibility may explain how GC persists despite coexisting with an aggressive invasive competitor, highlighting that competitive success is determined not only by winning aggressive interactions but also by maintaining access to essential resources (Boon et al., 2023; Alatalo and Moreno., 1987).

Our findings may also have important implications for understanding the impacts of invasive species on native freshwater fishes. Although GC exhibited considerable behavioural flexibility, the competitive efficiency seems to be poorer for the younger class, GC6. For example, though GC8 and GC11 did not significantly reduce their feeding in interspecific trials when compared to intraspecific ones, there was a significant reduction in feeding (with several trials having zero values) observed in the youngest developmental stage of GC (GC6) during interspecific contests, raising concerns about the survivability of the vulnerable ‘recruit’ stage of the species. In conclusion, our findings highlight that the behavioural plasticity exhibited by GC—particularly its capacity to modulate aggression levels and balance defensive hiding with active feeding—is a fundamental mechanism of resilience. By deciphering how native species navigate competitive pressures from invasive organisms, this study provides vital insights into the behavioural underpinnings of species persistence, offering a crucial framework for targeted conservation and management efforts in invaded ecosystems.

## Data Availability

All the raw data and R scripts used to analyse the data are available in Supplementary Information 2.

## Declaration of generative AI and AI-assisted technologies in the manuscript preparation process

During the preparation of this work, the author(s) used Google Gemini and Grammarly for checking any grammatical errors. The author(s) reviewed and edited the output as needed and take full responsibility for the content of the published article.

## Author contributions

VP was responsible for conceptualization, data curation, formal analysis, investigation, methodology, validation, visualization, and writing-original draft. PK was responsible for conceptualization, data curation, investigation, and methodology. RG contributed to conceptualization, formal analysis, funding acquisition, methodology, project administration, resources, supervision, writing - original draft, writing - review & editing.

## Acknowledgements

We are grateful to Chena Desai for assistance with fish maintenance.

