## Supplementary material for "Pick your battles: competitive strategies of green chromides (*Etroplus suratensis)* during inter- and intraspecific contests"

**Section 1. Supplementary Video Legends**

**Video S1.** Example of three bite events between an actor and a receiver. The behaviour is seen at 00:00, 00:02, and 00:03.

**Video S1 still:**

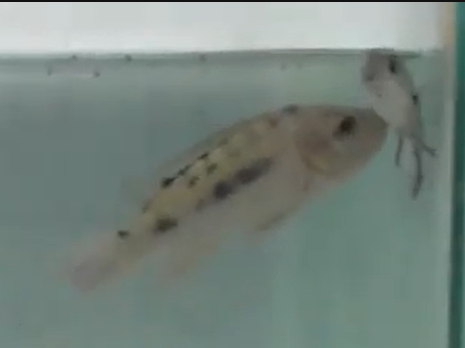

**Video S2.** Example of a charge event occurring at 00:01.

**Video S2 still:**

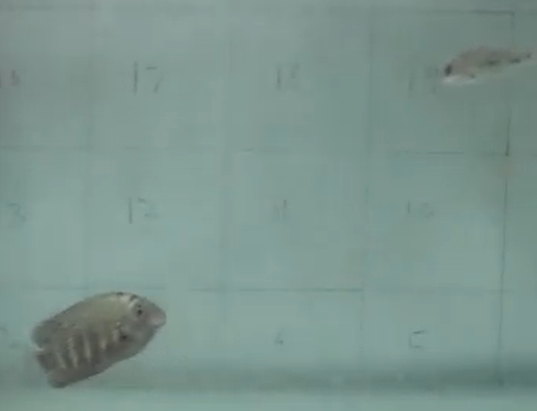

**Video S3.** Example of a chase behaviour, initiated at 00:00 and ending at 00:07.

**Video S3 still:**

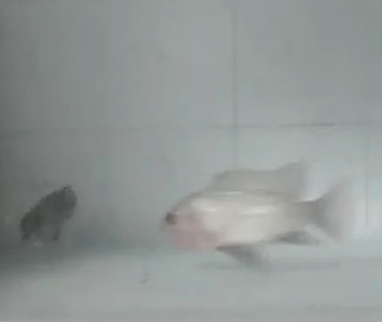

**Video S4**. Example of feeding bouts. The behaviour is seen at 00:01 and 00:05.

**Video S4 still:**

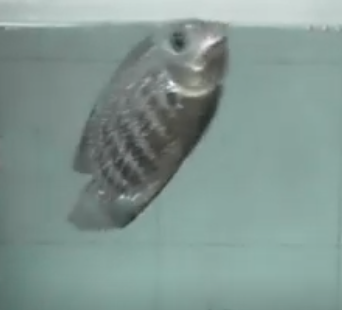

**Video S5.** Example of a hiding state, initiating at 00:01 and continuing past the end of the video clip, which is 00:03.

**Video S5 still:**

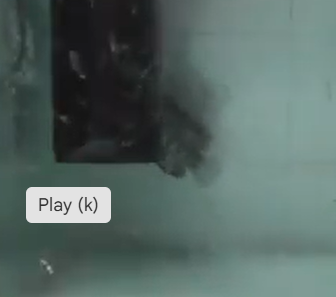

**Section 2. Supplementary Tables**

**Table S1.** Mean and range of behavioural responses exhibited by GC and T as R and I across all trial types.

| Trial type | GC | | | | T | | | |
| --- | --- | --- | --- | --- | --- | --- | --- | --- |
|  | R | | I | | R | | I | |
|  | Mean | Range | Mean | Range | Mean | Range | Mean | Range |
| **Aggression** |  |  |  |  |  |  |  |  |
| GC6_T6 | 9.67 | 0-43 | 2.25 | 0-9 | 0.58 | 0-2 | 2.08 | 0-10 |
| GC8_T6 | 3.83 | 0-13 | 2.42 | 0-11 | 1.92 | 0-14 | 0.5 | 0-2 |
| GC11_T6 | 7.45 | 1-19 | 10.5 | 4-20 | 2.55 | 0-7 | 1.91 | 0-4 |
| GC6_T8 | 3.58 | 0-12 | 4.83 | 0-40 | 14.2 | 0-41 | 2.17 | 0-11 |
| GC8_T8 | 21.8 | 8-60 | 13.3 | 5-31 | 7.08 | 1-14 | 4.5 | 0-14 |
| GC11_T8 | 9.58 | 1-20 | 10.1 | 1-28 | 3.67 | 0-9 | 1.08 | 0-3 |
| GC8_T11 | 1.67 | 0-10 | 0.08 | 0-1 | 135 | 7-274 | 19.7 | 0-78 |
| GC11_T11 | 6.88 | 0-29 | 1.19 | 0-10 | 91.3 | 0-316 | 19.2 | 0-131 |
| **Feeding** |  |  |  |  |  |  |  |  |
| GC6_T6 | 19.08 | 4-46 | 8.08 | 0-21 | 44 | 14-86 | 41.58 | 23-66 |
| GC8_T6 | 46.75 | 0-118 | 25.8 | 0-45 | 44.75 | 0-110 | 59.75 | 0-106 |
| GC11_T6 | 106.5 | 23-194 | 70.7 | 34-136 | 24.55 | 11-56 | 24.45 | 13-40 |
| GC6_T8 | 27.5 | 15-62 | 20.9 | 0-44 | 49.33 | 22-112 | 42.42 | 15-88 |
| GC8_T8 | 106.5 | 20-368 | 184 | 48-415 | 77.58 | 26-169 | 73.33 | 42-117 |
| GC11_T8 | 95.83 | 24-177 | 75.6 | 15-183 | 66.83 | 30-179 | 45 | 12-136 |
| GC8_T11 | 43.25 | 0-104 | 2.17 | 0-16 | 37.42 | 0-75 | 20 | 0-38 |
| GC11_T11 | 43.13 | 0-92 | 8.25 | 0-46 | 51 | 18-90 | 42.13 | 11-91 |
| **Hiding** |  |  |  |  |  |  |  |  |
| GC6_T6 | 1265.8 | 227-1717 | 1619 | 1241-1778 | 13.75 | 0-63 | 6.917 | 0-51 |
| GC8_T6 | 276.4 | 0-1147 | 99.5 | 0-604 | 32.67 | 0-155 | 25.17 | 0-69 |
| GC11_T6 | 68.55 | 0-615 | 3.73 | 0-32 | 90.64 | 0-670 | 127.5 | 5-319 |
| GC6_T8 | 923.6 | 0-1518 | 891 | 21-1800 | 140.4 | 0-607 | 2 | 0-11 |
| GC8_T8 | 237.3 | 0-761 | 22.1 | 0-97 | 45.83 | 0-227 | 31.67 | 0-135 |
| GC11_T8 | 219.6 | 0-901 | 18.1 | 0-141 | 47.5 | 0-293 | 101.1 | 0-231 |
| GC8_T11 | 699.3 | 0-1772 | 929 | 23-1819 | 24 | 0-67 | 3.167 | 0-15 |
| GC11_T11 | 185.6 | 0-1796 | 440 | 0-1776 | 0.938 | 0-6 | 0.125 | 0-2 |

**Table S2.** Estimates of the intraspecies GLMM model for total aggression

| Variable | Estimate | Std. Error | z value | Pr(>\|z\|) |
| --- | --- | --- | --- | --- |
| **Aggression: Negative binomial GLMM** |  |  |  |  |
| Intercept | 3.88085 | 0.3838 | 10.11164 | **4.91E-24** |
| Developmental stage MM | -0.64879 | 0.495886 | -1.30835 | 0.190756 |
| Developmental stage SS | -1.0446 | 0.348205 | -2.99995 | **0.0027** |
| Residency Status I | -0.70522 | 0.282799 | -2.49372 | **0.012641** |
| Contrast M | -0.68523 | 0.467483 | -1.46578 | 0.142709 |
| Contrast S | 0.399056 | 0.528108 | 0.755632 | 0.44987 |
| Contrast N | 0.806054 | 0.404226 | 1.994066 | **0.046145** |

P < 0.05 is indicated in bold

**Table S3.** Estimates of the intraspecies GLMM model for feeding bouts

| Variable | Estimate | Std. Error | z value | Pr(>\|z\|) |
| --- | --- | --- | --- | --- |
| **Feeding: Negative binomial GLMM** |  |  |  |  |
| Intercept | 4.557511 | 0.205655 | 22.16097 | **8.2E-109** |
| Developmental stage MM | -1.01819 | 0.199692 | -5.0988 | **3.42E-07** |
| Developmental stage SS | -0.97337 | 0.164178 | -5.92875 | **3.05E-09** |
| Residency Status I | -0.02349 | 0.10868 | -0.21611 | 0.8289 |
| Contrast M | -0.48287 | 0.270549 | -1.78477 | 0.074299 |
| Contrast S | 0.172215 | 0.266973 | 0.645066 | 0.518884 |
| Contrast N | -0.2143 | 0.234341 | -0.91449 | 0.36046 |

P < 0.05 is indicated in bold

**Table S4.**  Estimates of the intraspecies GLMM models for hiding

| Variable | Estimate | Std. Error | z value | Pr(>\|z\|) |
| --- | --- | --- | --- | --- |
| **Hiding occurrence: Binomial GLMM** |  |  |  |  |
| Intercept | -0.82885 | 1.064866 | -0.77836 | 0.436356 |
| Developmental stage MM | 8.460426 | 1.82463 | 4.636789 | **3.54E-06** |
| Developmental stage SS | 4.325667 | 0.960802 | 4.50214 | **6.73E-06** |
| Residency Status I | -1.03852 | 0.712106 | -1.45838 | 0.144735 |
| Contrast M | -6.16503 | 2.134772 | -2.88791 | **0.003878** |
| Contrast S | -1.49098 | 1.50806 | -0.98868 | 0.322821 |
| Contrast N | -0.947 | 1.434629 | -0.6601 | 0.509187 |
| **Hiding duration: Gamma GLMM** |  |  |  |  |
| Intercept | 5.46995 | 0.707288 | 7.733697 | **1.04E-14** |
| Developmental stage MM | 0.519636 | 0.721585 | 0.720131 | 0.471444 |
| Developmental stage SS | -0.07014 | 0.612205 | -0.11457 | 0.908782 |
| Residency Status I | 0.001232 | 0.267281 | 0.004608 | 0.996323 |
| Contrast M | 0.153344 | 0.901196 | 0.170156 | 0.864887 |
| Contrast S | -0.93012 | 0.745164 | -1.24821 | 0.211953 |
| Contrast N | -0.14572 | 0.691311 | -0.21079 | 0.833051 |

P < 0.05 is indicated in bold
